# Unraveling the viral dark matter of the rumen microbiome with a new global virome database

**DOI:** 10.1101/2022.11.30.518432

**Authors:** Ming Yan, Akbar Adjie Pratama, Zongjun Li, Yu Jiang, Matthew B. Sullivan, Zhongtang Yu

## Abstract

Like in the human gut and other environments, viruses are probably also diverse and modulate the microbiome (both population and function) in the rumen of ruminants, but it remains largely unknown. Here we mined 975 published rumen metagenomes for viral sequences, created the first rumen virome database (RVD), and perform ecogenomic meta-analyses of these data. This identified 397,180 species-level viral operational taxonomic units (vOTUs) and allowed for a 10-fold increase in classification rate of rumen viral sequences compared with other databases. Most of the classified vOTUs belong to the order *Caudovirales*, but distinct from those in the human gut. Rumen viruses likely have ecosystem impacts as they were predicted to infect dominant fiber degraders and methane producers, and they carry diverse auxiliary metabolic genes and antibiotic resistance genes. Together, the RVD database and these findings provide a baseline framework for future research on how viruses may impact the rumen ecosystem.

## Introduction

Recent virus-focused metagenomic studies of different habitats (ocean^1–3^, human gut^4–6^, soil^7^, and others^8,9,10^) have generated growing catalogs of virus genomes^4,5,11^, identified numerous auxiliary metabolic genes (AMGs)^12^, and revealed the vast diversity, community structure, and ecological impact of viruses^13^. Further, model system-focused efforts are beginning to map out the specifics of how viruses can metabolically reprogram their prokaryotic hosts in ways that lead to distinct ‘virocell’ states that alter the ecological fitness and outputs of their hosts^14^. At the ecosystem level, viruses are thought to have drastic impacts on ocean biogeochemistry^2,15,16^, human physiology^6^ and disease states^11^.

The rumen harbors a diverse multi-kingdom microbiome consisting of bacteria, archaea, fungi, protozoa, and viruses. Digesting and fermenting otherwise indigestible feedstuffs, the rumen microbiome provides ~70% of the energy (as volatile fatty acids)^17^ and ~80% of metabolizable nitrogen (as microbial protein) needed by ruminants to grow and produce meat and milk^18^. Recent metagenomic studies reveal strong associations of the rumen bacteria, archaea, and protozoa with feed efficiency, methane emissions, nitrogen excretion, milk and meat quality, and animal health^19^. However, rumen viruses, despite being abundant (5 × 10^7^ – 1.4 × 10^10^ virions/ml of rumen fluid^20^), were largely ignored. As obligate intracellular predators, rumen viruses can lyse their host cells and thus directly contribute to intra-ruminal recycling of microbial protein^21^, which decreases microbial protein (the major protein source for ruminants) outflow to the intestines and thus nitrogen utilization efficiency^22^. By altering the metabolism, ecological fitness, population dynamics, and evolution of their hosts, rumen viruses could conceivably affect the key functions and processes in the rumen.

While it has been challenging to study viruses in any complex system since most viruses cannot be cultured and they lack universal maker genes for detection and analyses^23^, virus-focused metagenomic approaches have powered several studies of rumen viruses. Using DNA from virion-enriched rumen samples, three studies identified viruses based on sequence similarity against reference databases^24–26^. They illuminated a diverse but small portion of the rumen virome because of the reference database limitations. Recent bioinformatics tools specific for viruses implementing machine learning algorithms (e.g., VirSorter2^27^ and VIBRANT^28^) and growing genomic resources (e.g., efam^29^) facilitate the identification of viruses from metagenomic sequences, and sequence-space organizational strategies provide scalable viral classification^30^ and taxonomy^31^. With these newly available bioinformatics tools and databases, two recent studies explored the diversity and ecogenomic implications of rumen viruses in 5 beef steers^32^ and one moose^33^. Although in both studies rumen viruses appeared tantalizingly important to the rumen ecosystem, only a few animals were involved. Here we seek to characterize the global rumen virome by (i) screening 20 TB of data from nearly 1,000 metagenomic datasets from diverse domesticated and wild ruminants across 5 continents, (ii) curating these data to establish a systematically cataloged rumen virome database (RVD), and (iii) examining the rumen virome in an ecogenomic context.

## Results

### The rumen viruses are highly diverse and represent unique lineages

Using state-of-the-art bioinformatics tools, we characterized the global diversity of the rumen virome by analyzing 975 published rumen metagenomes (Table S1) from 13 ruminant species (both wild and domesticated) under different husbandry regimes, across 8 countries and 5 continents (Fig. 1a and 1b). Following the recommendations of a recent viromics benchmarking paper^34^ and stringent criteria, we identified 705,380 putative viral contigs > 5 kb each and clustered them into 411,125 species-level viral operational units (vOTU) at > 95% of average nucleotide identity (ANI) and > 85% of coverage as recommended previously ^30^. We obtained a rumen virome database (RVD, download available at https://zenodo.org/record/7258071#.Y1ryEnbMK5c) representing 397,180 bona fide vOTUs (Fig. 1c), with 193,327 vOTUs being >10 kb. Checking with CheckV^35^ revealed 4,400 vOTUs being complete, 8,796 vOTUs > 90% complete, and 41,738 vOTUs > 50% complete. We assessed if RVD could improve identification of viral sequences across 240 rumen metagenomes^36^ over current virome databases. In total, RVD allowed for identifying around 3% of the metagenomic reads as viral sequences, which is >10 times higher than the mapping rate using IMG/VR^37^ (with the host-associated viruses only) while none of the reads could map to RefSeq Viral (Fig S1), demonstrating that RVD could greatly improve future rumen virome studies. Moreover, we noticed that subacute rumen acidosis (SARA), which is common among feedlot cattle and dairy cows fed a starch-rich diet, could significantly increase viral sequences in rumen metagenomes, consistent with profound SARA-associated variations of the rumen microbiome^38^.

**Fig. 1:**
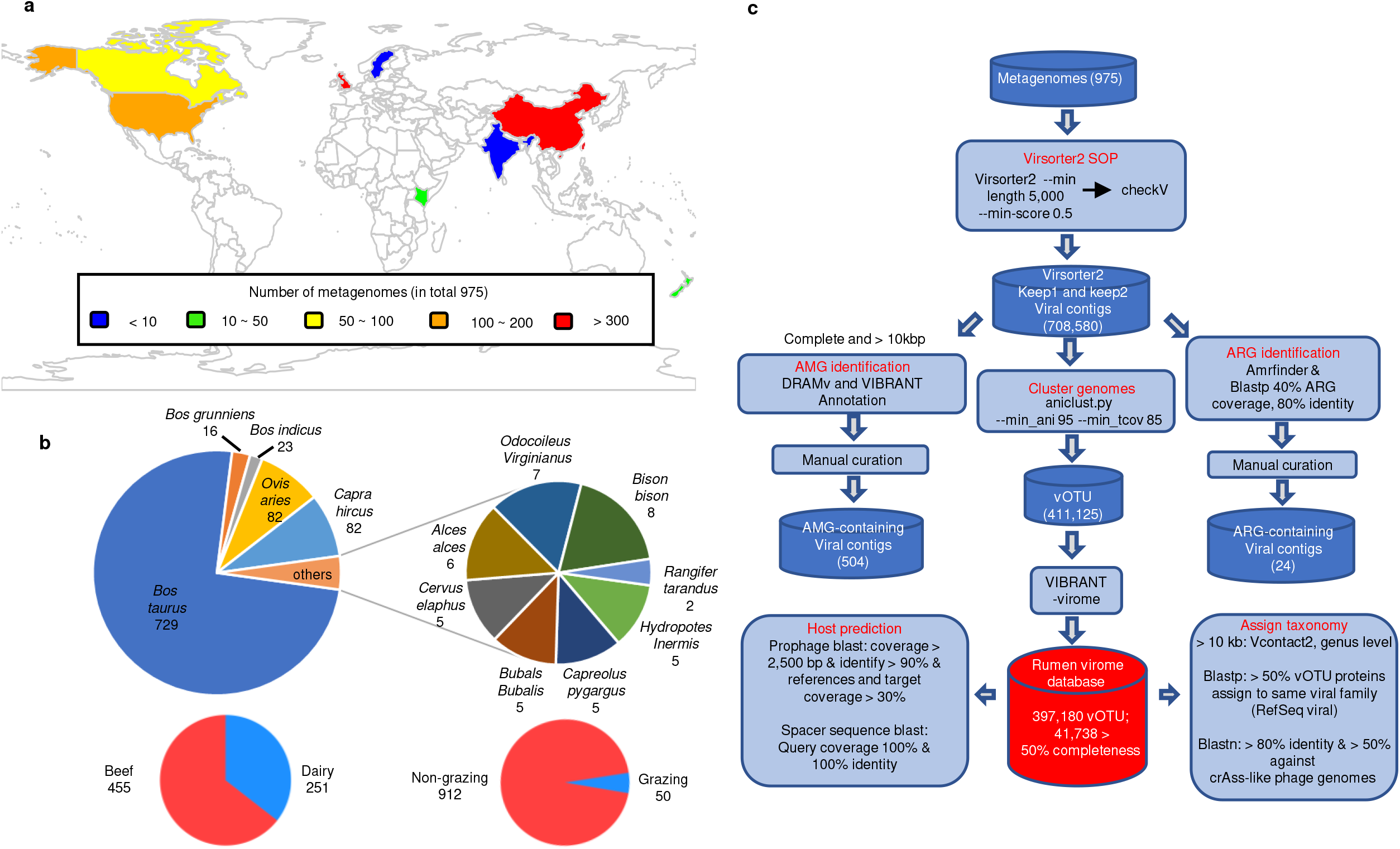
Rumen viral genomes recovered from 13 ruminant species across 5 continents. **a,** A global heatmap showing the geographic distribution of the 975 metagenomes and the proportion of the animals based on production system. **b,** Number of metagenomes from different ruminant species or production husbandries. **c,** Workflow of the rumen virome analysis pipeline. See also Table S1 for detailed information about the metadata.

We classified 1,857 (0.47 %) of the vOTU to existing genera using vConTACT2^31^ and another 32,801 vOTUs (8.3% of the total) only to an existing family. Most of the family-level vOTUs (98.4%) were assigned to the families *Siphoviridae, Myoviridae, Podoviridae* in the order *Caudovirales* (Fig. 2a, 2b.). These three families were also identified as the most abundant classifiable viruses in the human gut viromes^4,39^. The order *Caudovirales* had most of the vOTUs (99.7%) taxonomically classifiable at the genus and family levels, as in the case of the human gut virome^4^. We compared all the genus-level vOTUs of *Caudovirales* between the human and the rumen viromes using a phylogenetic tree (Fig. 2c). Only 14% of the *Caudivirales* genera were shared between the two types of viromes (Fig. 2d), demonstrating their divergence. The remaining vOTUs (91.7% of the total) could not be assigned to any existing taxa, indicating that most of the rumen viruses represent new lineages and are colossally underrepresented in the current virome databases. Additionally, 121 vOTUs (0.03%) were identified as crAss-like viruses.

**Fig. 2:**
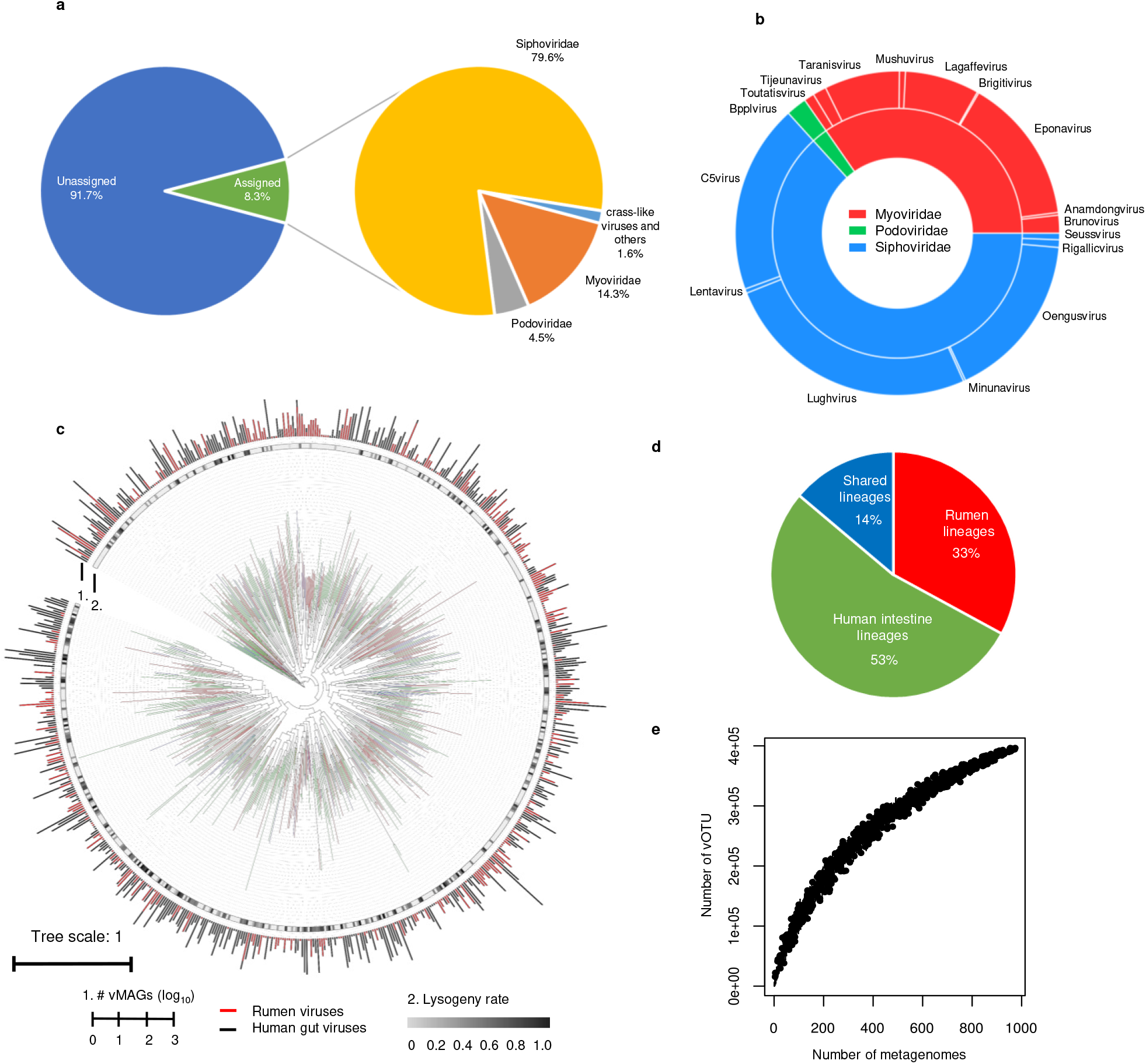
Taxonomic classifications of the rumen viruses. **a,** Family level taxonomy and its proportion of the rumen viruses in the rumen virome database RVD). Most of the vOTUs (99.7%) classified to existing genera or families were under the order of *Caudovirales*, including 121 identified crAss-like phages. Detailed taxonomy assignment for individual vOTUs could be found in Table S2. **b,** Genus-level taxonomy and proportion of the 1,858 vOTU assigned using vConTACT2. **c,** A phylogenetic tree of *Caudovirales* viruses built with 77 concatenated marker genes identified in 10,203 viral metagenome-assembled genomes (vMAGs) with a > 50% completeness of the current study and the two largest human gut virome databases (MGV^1^ and GPD^2^). For better visualization, only one representative vMAG (the longest and most complete) per genus-level vOTU was included (in total 714 genomes). The branches were color coded: green, the *Caudovirales* lineages exclusively from the human virome; red, the lineages exclusively found in the rumen virome of the current study; blue, the lineages found in both the rumen and the human viromes. Lysogeny rates were calculated with VIBRANT and shown as the inner ring. The number of vMAGs representing each lineage was shown as barplot (red for human viruses, and black for human viruses). **d,** Proportion of lineages *Caudovirales* viruses between the rumen and the human viromes. **e,** A rarefaction curve of the vOTUs identified in the rumen virome. The upward trend of the rarefaction curve indicates that more rumen viruses remain to be identified at the specie level.

We also identified eukaryotic viruses, some of which were assigned to the families *Phycodnaviridae, Mimiviridae*, and *Retroviridae*. In contrast to the human gut microbiome, which has few eukaryotes, the rumen microbiome has diverse fungi and protozoa making up to 50% of the microbial biomass thereof and they play important roles in fiber digestion and nitrogen recycling, respectively^40^. The revelation of rumen eukaryotic viruses is thus not unexpected. Based on host prediction, viruses infecting archaea were also identified. Although we focused on dsDNA viruses, we identified 109 vOTUs as ssDNA rumen viruses, all assigned to the family *Microviridae*. Add more info on the type(s) of microbes they infect. Given the genome size cutoff (5 kb) we used, ssDNA viruses, which have a relatively smaller genome, might be underrepresented in RVD. Indeed, ssDNA viruses are enriched in viral metagenome^4^, and hence future research focusing on ssDNA viruses should use virion-enriched metagenomes.

Rarefaction analysis (Fig. 2e) revealed a diverse global rumen virome, and its full diversity remains to be revealed. Additionally, this study only analyzed DNA viruses. Because RNA viruses are also likely diverse, abundant, and of great ecological importance, as demonstrated in the ocean^1,15^, future studies should include rumen RNA viruses once more metatranscriptomic datasets are available.

### Rumen viruses have a broad range of hosts compared with human gut viruses

We predicted the hosts of the rumen prokaryotic viruses by matching the spacer sequences and prophage sequences, with conservative thresholds, with 251,167 reference gnomes of prokaryotes in NCBI RefSeq (Release 211) plus 25,234 metagenome-assembled genomes (MAGs) of rumen prokaryotes (see Methods). Ciliate hosts were inferred using 52 singe-cell amplified genomes (SAGs) of rumen ciliates^41^. We did not predict the hosts of fungal viruses due to the lack of genomes of rumen fungi. In total, species of 25 genera of archaea and 1,051 genera of bacteria were predicted to be likely infected by the identified phages. We generated a genome-based genus-level phylogenetic tree of the bacterial hosts (Fig. 3a) and the archaeal hosts (Fig. S2) with their predicted phages to examine the lysogeny rate, number of phages per genus of hosts, and number of phages per genome. Similarly, we also generated a genome-based phylogenetic tree of the 52 SAGs and their phages and estimated the number of phages carried by individual SAGs (Fig. S3). Out of the 52 ciliate SAGs representing 19 species across 13 genera, 38 (including all the species) had predicted prophage sequences, suggesting a high prevalence of ruminal phages potentially infecting ciliates.

**Fig. 3:**
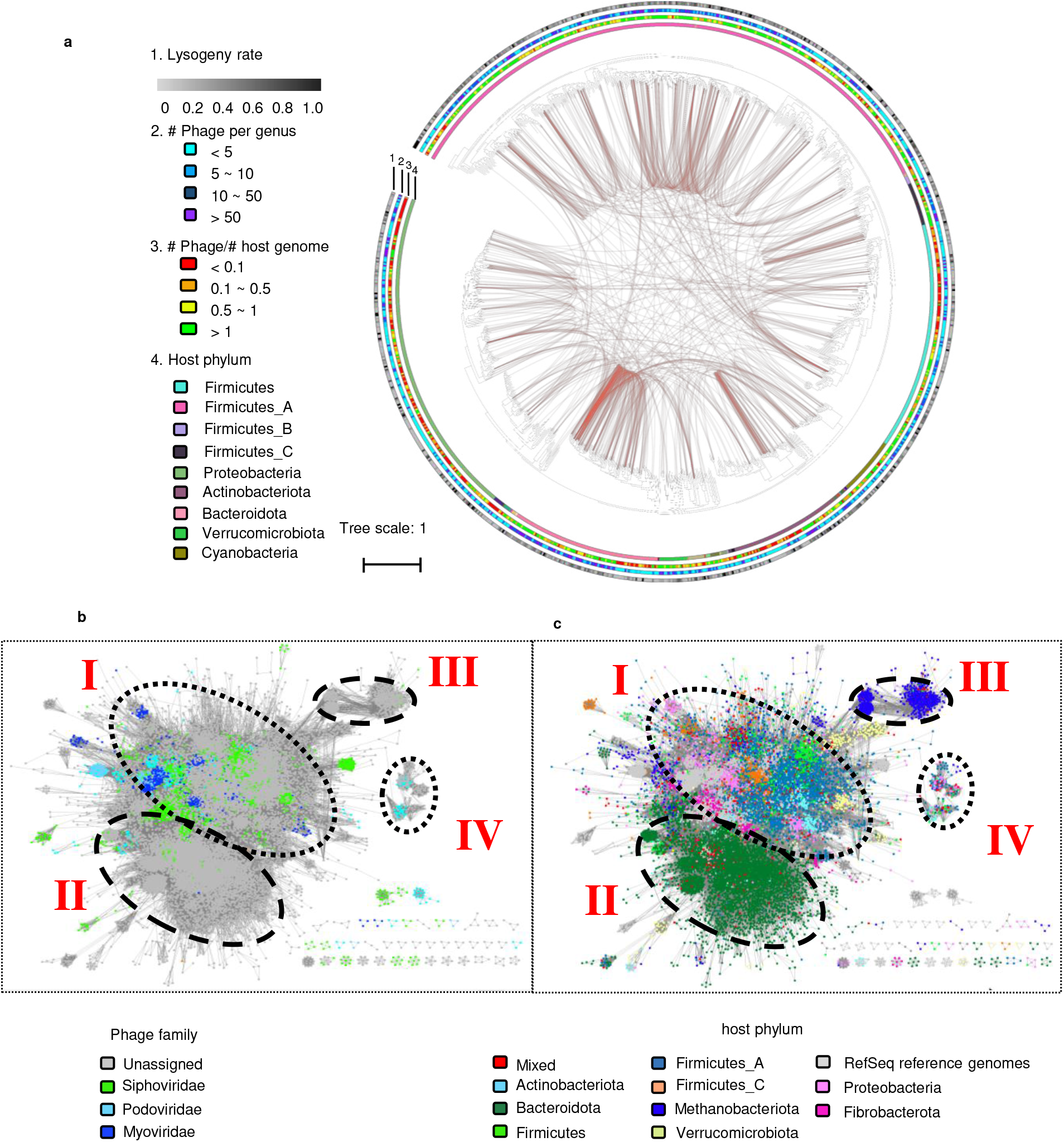
Bacterial host range of the rumen viruses. **a**, A genome-based phylogenetic tree of 1,051 bacterial genera that contained the predicted hosts of 40,881 vOTUs. The hosts were inferred by (i) aligning the sequences of the representative vMAGs (the longest with the highest completeness) of each vOTUs with 22,087 metagenome-assembled genomes (MAGs) of rumen bacteria and 242,387 bacteria reference genomes of NCBI RefSeq and (ii) aligning the CRISPR spacer sequences of the representative vMAGs and those of the prokaryotic genomes and MAGs. The prokaryotic genomes and MAGs were classified using GTDB-Tk. The phylogenetic tree was constructed with the genomes or MAGs of the inferred hosts (clustered into genera) and their predicted phages to examine the lysogeny rate, number of phages per genus, and number of phages per genome/MAG. In total, 4,394 vOTUs likely had a host range across multiple genera. These genera are connected with orange arcs. The rings correspond to lysogeny rates (ring 1, calculated based on VIBRANT results), number of phages per genus of hosts (ring 2), number of phages per genome/MAG (ring 3), and bacterial phyla to which the bacterial host genera belon (ring 4). **b,** Gene-sharing network of the vOTU with predicted host obtained from vConTACT2 and visualized using Cytoscape. The virus genomes were colored according to the family assignment (left) and predicted host phylum (right) respectively. See also Figure S1 and Figure S2 for the host assignment of archaea and protozoa and Table S3 for the detailed host assignments for individual vOTU.

More vOTUs were predicted to be bacteriophages than archaeophages (40,881 vs. 2,403). Phages can infect multiple host species^5,11,42^, and 9,214 (22.5%) bacteriophages and 396 (16.5%) archaeophages of the rumen virome could infect multiple host species. The proportion of rumen bacteriophages with a broad-host range is lower than that of human gut bacteriophages (about a third^5,42^). The differential may be explained by the higher coverage of phages in the human gut metagenome databases. However, we found that 3.8% (1,544) of the broad-host-range rumen phages could potentially infect species across multiple bacterial phyla (Fig. 3a). Even excluding the host species inferred from the RefSeq genomes of prokaryotes, which contain genomes generated from various ecosystems (including the rumen), we still found that 3.5% of the broad-host-range phages potentially infect multiple phyla of rumen bacteria. This rate is much higher than that (merely 0.13%) observed in the human gut microbiome^5^. This differential may be attributable to a more diverse microbiome in the rumen than in the human gut. Indeed, we found that all 32 phyla of the reference rumen bacterial genomes could be infected by phages, while only 8 phyla were likely the host of human gut phages. In addition, rumen digesta is more homologous, due to high fluidity and continuous mixing, than the human gut digesta. The rumen microbiome may have more gene flows among microbes due to mixing and physical proximity than the human gut microbiome. The broad host range of rumen phages reveals their potential in mediating gene flow across phylogenetic boundaries of bacteria, which can facilitate microbial evolution and adaptation^5,42^.

The theory of “local adaptation” was proposed based on the finding that sympatric phages are more infective than allopatric phages in soil^43^. Similarly, sequence identity analysis of the human gut microbiome revealed that homologous phages usually infect homologous MAGs^42^. However, analysis of sequence identity failed to classify viruses with variable evolutionary mode and tempo^31^. To better examine the relationship between phages and their hosts and the gene flow, we generated a gene-sharing network of all the phages with their taxonomy (Fig. 3b) and their host phyla (Fig. 3c). From the 43,284 vOTUs with a host match, we found 2,764 genus-level clusters, of which only 218 clusters included one or more RefSeq viral genomes. Therefore, RVD greatly expanded the RefSeq viral genomes (by > 12-fold) at the genus level.

The gene-sharing network grouped most of the rumen vOTUs with their predicted hosts into four groups (Fig. 3b). Groups I (the largest) and IV (the smallest) had more classified vOTUs than groups II and III. Groups II and III mainly infected *Bacteroidota* and *Methanobacteriota*, respectively, while groups I and IV had a broader range of phyla (Fig. 3c). *Bacteroidota* species can degrade and utilize major feed ingredients (hemicellulose, starch, and protein), and it is the most abundant phylum of the rumen microbiome^44^. As the most predominant genus in the rumen, *Prevotella* accounts for 40-60% of the total copies of bacterial 16S rRNA gene^45^, while *Methanobacteriota* is dominated by the genus *Methanobrevibacter*, which can reach >63% of the total methanogens^46^. The narrow host range (a single phylum) of groups II and IV corroborates that phages with a high degree of gene-sharing generally have a homologous host range. Most of the vOTUs inferred to infect *Bacteriodota* and *Methanobacteriota* could not be classified with the current virome databases, suggesting that they represent new viral lineages. Groups I and IV were predicted to infect multiple phyla of bacteria, including both Gram-positive and Gram-negative bacteria that have different niches and capacities, but none of their genera or families are predominant in the rumen. Some small clusters were evident within groups I and IV. They might represent individual phyla or lower taxa (e.g., genera) of phage hosts. Thus, the theory of “local adaptation” may explain the host range of the phages in these two groups. It remains to be determined if the greater extent of metabolic interactions and functional redundancy in the rumen than in the human gut explain the different extents of local adaptation of viruses.

### Some rumen viruses may ameliorate the fitness and ecological impact of their hosts by providing auxiliary metabolic genes (AMGs)

Phages can alter host metabolism and physiology by forming virocells upon infection^47^. One underpinning mechanism is to provide hosts with AMGs, which can potentially impact ecological processes, including global carbon recycling^48^ and nitrogen metabolism in the ocean^49^, sulfur metabolism in the environment^12^, and organosulfur metabolism in the environment and the human gut^50^. In compassion, AMGs carried by rumen viruses have only been reported in one study that involved 5 beef steers^32^. We thus sought to identify the AMGs encoded by the rumen virome. To be conservative, we only searched the complete viral MAGs (vMAGs) for AMGs after a series of cautious annotations and manual curation (see Methods). In total 504 complete vMAGs were found to carry one or more AMGs (see Table S4 for the annotation and the curation results), which represent 62 different AMGs (Fig. 4b), including 22 that were identified in at least two vMAGs of RVD and 49 that have been identified in previous studies. These AMGs were involved in different types of metabolism (Fig. 4a), including carbohydrate utilization, nitrogen metabolism, nucleotide metabolism, signaling, transport, and others. Given our stringent curations, the large number of incomplete vMAGs, and viruses that remain to be identified, the rumen virome likely carries more AMGs.

**Fig 4:**
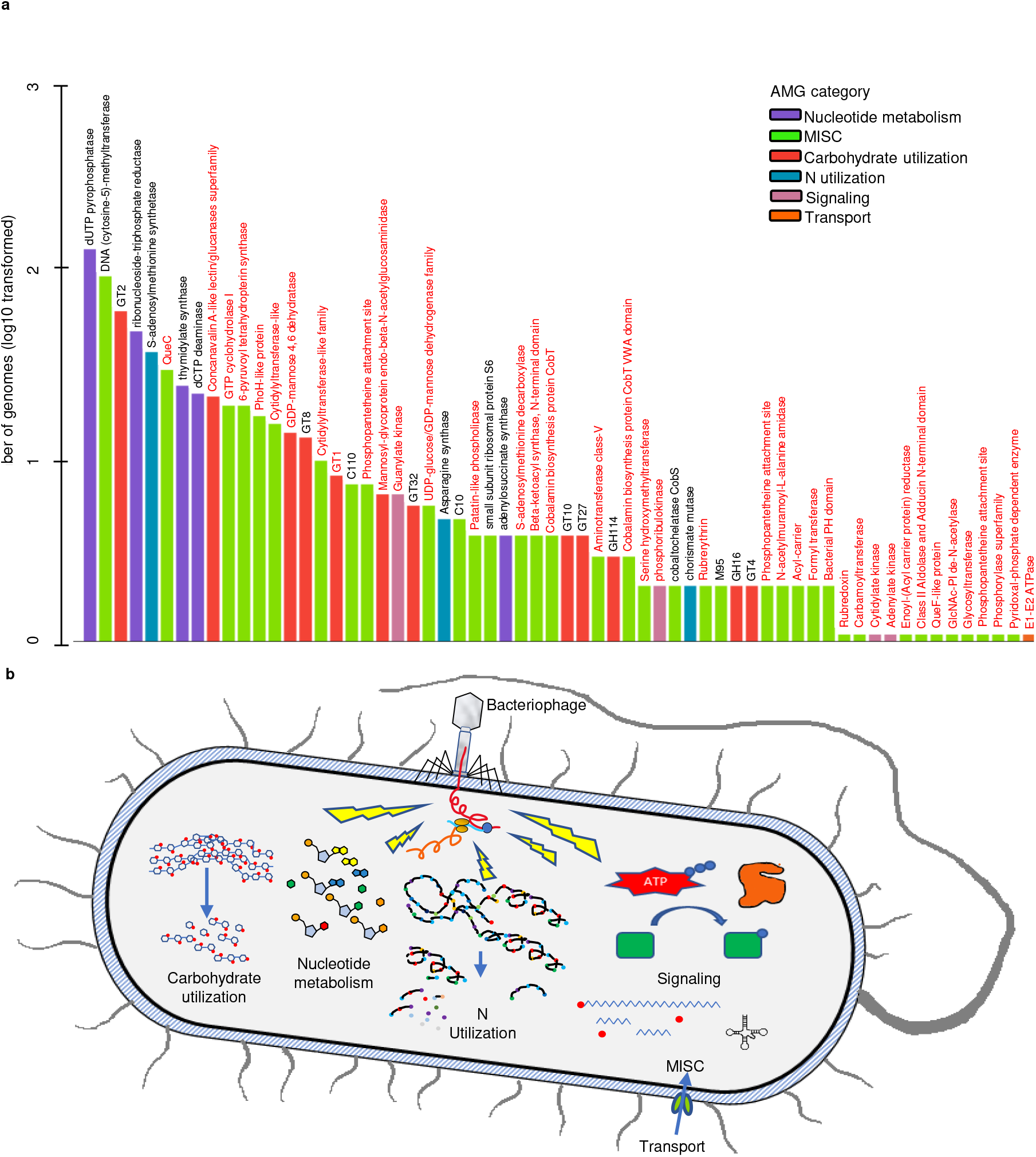
Auxiliary metabolic genes carried by rumen viruses (a) and a conceptual model illustrating how AMGs might enable rumen viruses to modulate host metabolism (b). AMGs were predicted only from complete vMAGs (504 in total) that passed a series of curation, and only the AMGs that had been identified in previous studies (labeled in red) and/or identified in more than two vMAGs of the current study were shown. See Table S4 for the detailed AMGs curation process, full annotation of the final AMG-carrying vMAGs, and AMG functional category annotation.

The AMGs were annotated to many categories of metabolism (Fig. 4b). More AMGs were involved in nucleotide metabolism than other metabolism, corroborating their important role in shifting host nucleotide metabolism to benefit viruses, which has been verified experimentally with two AMGs: thymidylate synthase gene (*thyA*) in marine viruses^14^ and dCTP deaminase gene (*dcd*) in *Bacillus subtilis* viruses^51^. We found DNMT1 as the second most prevalent AMG in nearly 100 rumen vMAGs. This is in line with the high prevalence of this AMG in ocean virome^52^. DNMT1 encodes DNA (cytosine-5)-methyltransferase that can help viruses evade the antiviral restriction-modification systems of their host^53,54^. Carrying an AMG that encodes an orphan DNA methyltransferase might be one defense mechanism among rumen viruses. Several identified AMGs encode carbohydrate-active enzymes (CAZymes), including glycoside hydrolases (GHs) and glycosyl transferases. Conceivably, the GH type of AMGs can enhance the ability of bacterial hosts to utilize polysaccharides. Asparagine synthase, a key enzyme in ammonia assimilation by rumen bacteria^55^, was also encoded by one of the identified AMGs. The AMGs carried by some ruminal viruses may not only enhance their survival but also greatly impact feed digestion, nitrogen metabolism, and other activities and hence ruminant production.

### Rumen viruses carry a few types of antibiotic resistance genes (ARG)s but may facilitate ARG transmission across phylogenetic boundaries

We searched ARGs among the 705,380 viral contigs using stringent criteria against two expert-curated databases: CARD^56^ and the NCBI’s Bacterial Antimicrobial Resistance Reference Gene Database^57^, and we found 24 contigs carrying ARGs. The dearth of ARG-carrying rumen viruses corroborates the previous finding that phage genomes rarely carry ARGs^58^. The rumen virome may not be a significant reservoir of ARGs, but it could still transmit ARGs among rumen microbes as several major ARG classes were found, including tet(W) and tet(O) (Fig. 5a and Fig. S4), both of which are prevalent and highly expressed in the rumen microbiome^59–62^. Three vMAGs were recovered from three different rumen metagenomes of the same herd, but they carried the same ARGs (Van(G) and Van(W-G)) and nearly identical genomic architecture (Fig. 5c), pointing to a potential of rumen viruses as a route of ARG transmission between different animals. Even with the strictest criteria requiring ARG flanking by two viral hallmark genes, we still found several vMAGs (Fig. 5c and Table. S5). A comparison of the prevalence of ARG-carrying viruses among different animal husbandry regimes revealed a higher prevalence of ARG-carrying viruses in beef cattle than in dairy cattle and in non-grazing animals than in grazing animals (Fig. 5b). These findings are not totally surprising because more antimicrobials are fed to beef cattle than dairy cows and non-grazing animals than grazing animals. Another possible explanation for the above differentials may be diets: beef cattle and non-grazing animals are fed a higher portion of concentrate (starch) than dairy cows and grazing animals, respectively; and a higher ARG prevalence correlates with high-concentrate diets^63^.

**Fig 5:**
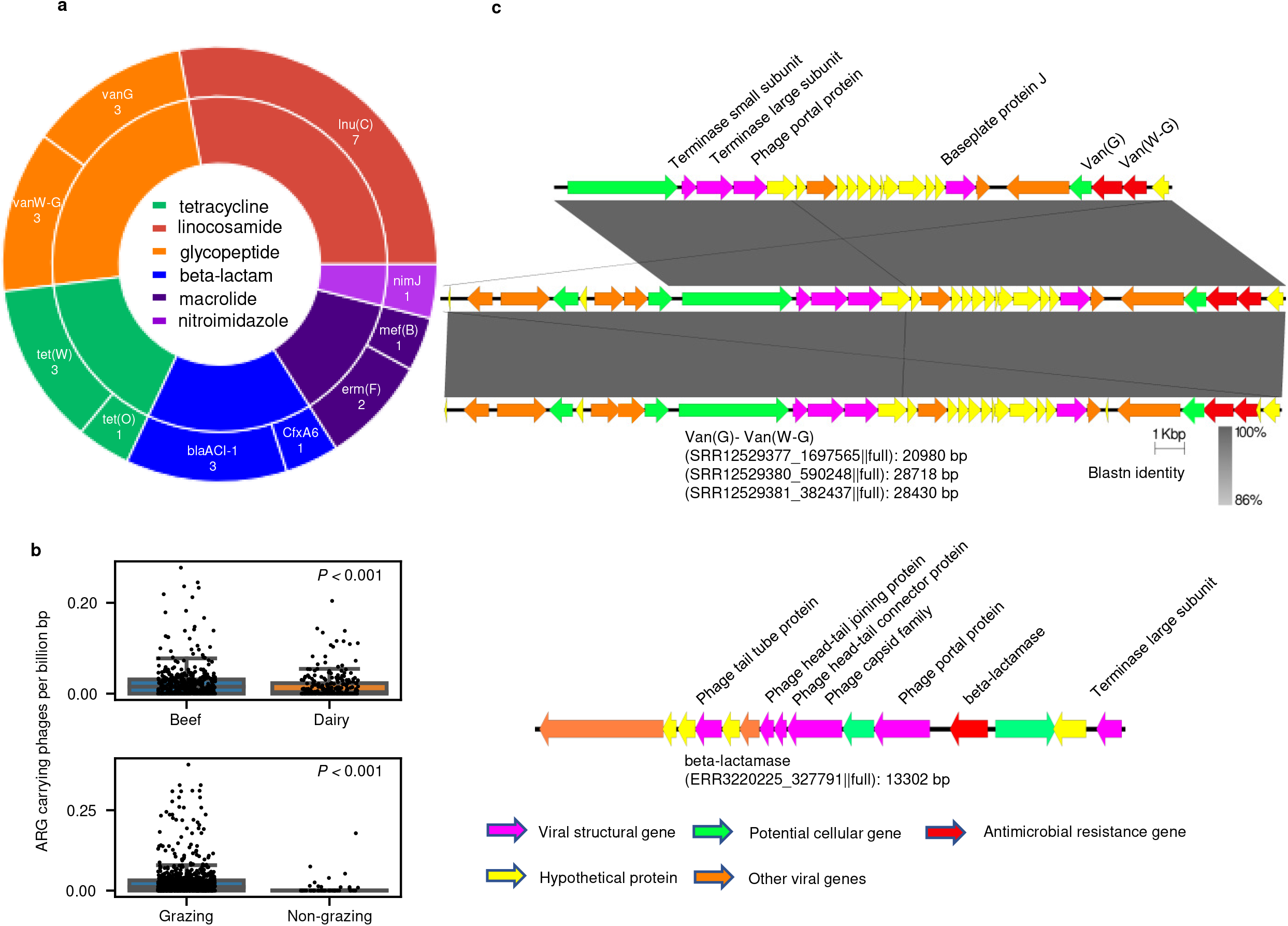
ARG carried by rumen viruses exhibiting distinctive abundance in different countries and in different animal husbandry paradigms. **a,** The number of ARG-carrying viruses and their ARG classes identified in the selected vial contigs (24 out of 705,380). **b,** The abundance of ARG-carrying viruses (number of ARG-carry viral contigs per Gb of metagenomic DNA sequences) in different countries and different animal husbandries (beef vs. dairy and non-grazing vs. grazing). Box plots showing the median and quartiles of the prevalence of ARG-carrying viruses. Statistical significance was tested using the non-parametric Wilcoxon signed-rank test. **c,** The genomic organization of three representative ARG-carrying viral contigs. These three viral contigs shared the same two *Van* genes with highly similar sequences and had nearly identical genomic organization but were found in three different samples. The lines connecting the gene among contigs indicating blastn identity. The contig on the bottom represents the viral contig with ARG flanked by viral hallmark gene. The representative viral contigs carrying other ARG genes were chosen based on CheckV completeness and the number of cellular genes, and their genome organizations are displayed in Fig S4. The manual curations of each ARG-carrying contigs and their detailed annotation could be found in Table S4.

### Rumen virome is highly individualized but a core virome exists within each species or type of ruminants

Motivated by the finding that crAss-like viruses are highly prevalent in the human gut^64^ and the axiom that a “core” virome may exist in the human gut microbiome^65^, we searched RVD for “core” rumen virome based on prevalence in different species or types of ruminants. We identified a “core” rumen virome (present in > 50% of the samples) in each species or type of animals (Fig. 6a). The “core” rumen virome only accounts for <0.01% of the total viral diversity, signifying highly individualized rumen virome (Fig. 6a). The majority of the “core” virome were taxonomically unassigned, but they were predicted to infect the “core” rumen bacteriome (Table S6). Unlike in the human gut virome, none of the rumen crAss-like viruses were included in the rumen “core” virome. The “core” rumen virome in each species or type of animals was estimated to account for around 10% of the total rumen virome (Fig. 6b), suggesting that it may have a great impact on regulating the population and functions of the rumen microbiome. Different species or types of ruminants might also have a different “core” rumen virome (Fig. 6c), likely reflecting the difference in the rumen microbiome among the species or types of ruminants.

**Fig 6:**
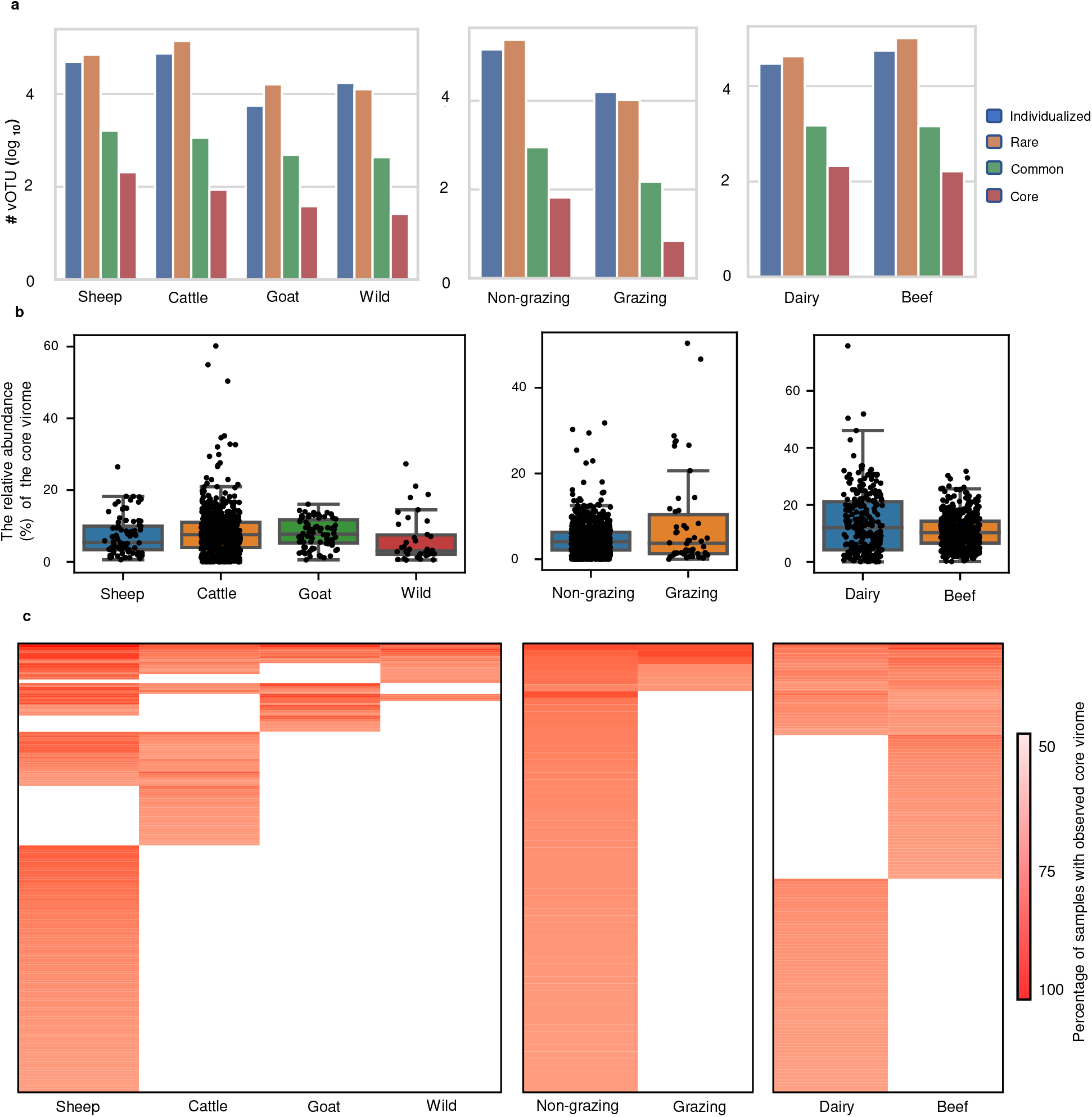
Rumen virome is individualized but a core virome population exists. **a,** The number (log10 transformed) of vOTUs observed in only one sample (Individualized), ≥2 samples but less than 20% of the animal group (Rare), 20 - 50% of the animal group (Common), and >50% of the animal group (Core). **b,** Heatmap of the occurrence of core virome in each animal group. Only the groups with >38 metagenome datasets were included. Detailed information of the core vOTU (family-level taxonomy and predicted hosts) could be found in Table S6. **c,** The relative abundance (%) of the core virome in each animal group. The number of samples in each group are 92 for sheep, 729 for cattle, 72 for goats, 38 for wild ruminants (including *Alces alces, Bison bison, Capreolus_pygargus, Cervus_elaphus, Hydropotes_inermis, Odocoileus_virginianus, Rangifer_tarandus*), 912 for non-grazing ruminants, 50 for grazing ruminants, 251 for dairy cows, and 455 for beef cattle.

## Discussion

Recent studies have documented the vast diversity and potential ecological impact of viruses in the environments (ocean, soil, anaerobic digester, etc.) and the human gut. However, the viruses residing in the rumen are poorly characterized and understood^66^. As the first comprehensive study on the rumen virome, we mined most of the published rumen metagenomes (nearly 1,000), recovered 397,180 species-level vOTUs, and created the first rumen virome database, RVD. This database greatly expanded the databases of rumen viruses and shined new lights on their diversity across major ruminant species, both domesticated and wild. Because nearly 92% of the identified vOTUs could not be classified to any existing taxa, most rumen viruses represent new lineages.

Using an integrated approach and conserved thresholds, we predicted that rumen viruses could infect diverse rumen microbes including those with great functional and environmental importance. Of the rumen vOTUs with a predicted host match, 99.5% of them were inferred to infect ruminal prokaryotes even though most of the reference genomes were from non-ruminal prokaryotes, demonstrating the rigorousness and low false positive rate of our host prediction pipeline. Surprisingly, rumen viruses seem to have a broader host range than those in other ecosystems as evidenced by the high proportion of rumen vOTUs predicted to infect multiple hosts across phylogenetical boundaries, even phylum boundaries. This is unlikely due to false prediction as isolated rumen viruses can indeed infect species of both *Bacteroidota* and *Firmicutes*^67^. The broad host ranges of ruminal viruses signify their important roles in facilitating host evolution.

We found diverse categories of AMGs, including those encoding enzymes modifying host nucleotide metabolism and providing antiviral defense, both of which can buttress viral fitness and have been demonstrated experimentally in other ecosystems^51,54^. Glycoside hydrolases are the most important enzymes of the rumen microbiome as they enable the digestion of plant-based feed therein. Thus, the revelation of AMGs encoding GHs, which was also reported in the rumen previously^32^, suggests that some rumen viruses may potentially enhance feed digestion by ruminants. It should be noted that we only predicted AMGs from a small portion (504 out of 397,180) of the rumen viruses identified. The rumen virome could carry orders of magnitude more AMGs. The other side of the coin is that lytic rumen viruses can aggravate intra-ruminal nitrogen recycling by lysing their host cells. Therefore, the rumen virome likely has a great impact on feed digestion and microbial protein yield through phage-host interactions, potentially affecting ruminant production.

We identified ARGs, including a few major classes of ARGs, carried by rumen viruses. One of the most prevalent ARGs, tet(W), identified in this study had been found in an integrative and conjugative element (ICE) recently^68^. Three vMAGs from three different rumen microbiomes from the same herd carried the same ARG and had nearly identical genomic architecture. These findings and the broad host range of ruminal phages highlight the potential of rumen viruses to transmit ARGs, especially in beef and non-grazing animals. It should be noted that although ARGs appeared to be sparse in the rumen virome, much of the viral diversity remains to be uncovered. More ARGs can be expected in future rumen virome studies.

Collectively, this study provides the largest rumen virus database (RVD) hitherto and new insights into the diversity and potential role of the rumen virome in regulating the metabolism, feed digestion, microbial protein synthesis and degradation by their cellular hosts. The RVD database will greatly facilitate future studies on the rumen virome. Furthermore, the finding of the current study can help inform future ecological studies to investigate how rumen viruses may regulate the rumen microbiome and its function, and thus ruminant production.

## Methods

### Assembly and identification of viral genomes from rumen metagenomes

Published rumen metagenomes (n=975, Table S1) were downloaded from NCBI SRA, quality controlled with fastp (v0.23.1)^69^, and then assembled individually using Megahit (v1.2.1) with the default parameters. The 975 metagenomes were published in 80 studies covering 13 species/breeds of ruminants from 8 countries (Table S1) across 5 continents (Fig. 1). The countries where the metagenomes were sampled and the number of metagenomes were visualized on a world map using the R package ‘rworldmap’.

After assembly, tentative viral contigs were first identified following the viral sequence identification SOP (dx.doi.org/10.17504/protocols.io.bwm5pc86). Briefly, tentative viral contigs > 5kb were further verified using VirSorter2^27^ (option: --min-score 0.5), and the resultant contigs were piped to CheckV^35^ to trim off host sequences flanking proviruses. We only chose viral contigs >5 kb because the currently available bioinformatics tools have a relatively high false positive rate when identifying viral contigs <5 kb ^34^. Only the contigs falling into categories Keep1 and Keep2 were retained as putative viral contigs (708,580 in total) for further analyses.

The viral contigs were clustered into vOTU at 95% ANI over 85% of the shortest contigs as suggested^30^ using a custom script from the CheckV repository^35^. The resultant 411,125 vOTUs were then further verified with VIBRANT ^28^ (option: --virome). We obtained 397,180 bona fide vOTUs (96.6%), which were identified to be of viral origin. To be conservative, only the vOTUs identified by both VIBRANT and VirSorter2 were used to build RVD and retained for taxonomy classification and host prediction. We chose to use both VIBRANT and VirSorter2 for viral identification because they are among the latest tools with the best performance according to a recent benchmarking paper^34^. We were interested in determining how viral reads could be annotated with the existing databases, but no rumen viral database was available. Thus, we annotated the clean reads using complete genomes from RefSeq viral (release 211; downloaded in March 2022) and the host-associated fraction of the latest IMG/VR database^37^. The completeness of the vOTUs was estimated using CheckV^35^. A rarefaction curve was generated to assess the coverage of rumen viruses using the package RTK^70^ in R.

### Taxonomic classification of all vOTUs and tree construction of those assigned to the order *Caudovirales*

We assigned vOTUs >10 kb to genus-level taxa based on a gene-sharing network using vConTACT2^31^, which uses RefSeq Viral (release 88) as reference genomes. The vOTUs that could be clustered with the reference genomes of a genus were assigned to that genus according to the vConTACT2 workflow. We assigned the vOTUs that failed to be assigned to a genus and those <10 kb to family-level taxa using the majority rule as done previously^6^. Briefly, we predicted the ORFs of each vOTU using Prodigal^71^ and then aligned the ORF sequences with those of RefSeq Viral using BLASTp with a bit score of ≥50. The vOTUs that aligned with the RefSeq genomes of a family with over >50% of their protein sequence were assigned to that family. We identified crAss-like phages using BLASTn against 2,478 crAss-like phage genomes identified from previous studies^64,72,73^ with the threshold of ≥80% sequence identity over ≥50% of the length of previously identified crAss-like vMAGs.

Most of the taxonomically assignable vOTUs of the rumen virome, as in the case of the human gut virome, were assigned to the order *Caudovirales*, and thus we compared the phylogenic distribution of the *Caudovirales* viruses between the rumen and the human gut viromes based on concatenated protein phylogeny^74^. Specifically, we downloaded the HMM profiles of the 77 marker genes of *Caudovirales* viruses from VOGDB (http://vogdb.org) and searched RVD of the current study and the two largest human gut virome databases (MGV^4^ and GPD^5^, which phylogenetically complement each other) for the marker genes using search HMMER v3.1b2^75^. To ensure a fair comparison across the databases, only the vMAGs with a completeness >50% were included in the search. We then aligned each of the marker genes from the three databases using MAFFT^76^, sliced out the positions where more than 50% were gaps using trimAl^77^, concatenated each aligned marker gene, and filled the gap where a marker gene was absent. Only the concatenated marker gene each with >3 marker genes and found in > 5% of all the aligned concatemers were retained, resulting in 10,203 *Caudovirales* marker gene concatemers each with 13,573 alignment columns. These marker gene concatemers were clustered into genus-level vOTUs as described previously^4^, which has been benchmarked to achieve high taxonomic homogeneity using viral RefSeq genomes. We built a phylogenetic tree of *Caudovirales* viruses using FastTree v.2.1.9 (option: -mlacc 2 -slownni -wag)^78^ and the alignment of the concatenated marker genes of the representative vMAG sequences of all the genus-level vOTUs with the highest genome completeness (> 50%, based on CheckV analysis). The *Caudovirales* tree was visualized using iTOL^79^. The vMAGs identified as prophages or encoding an integrase were considered lysogenic, and lysogenetic rate was calculated based on the VIBRANT results.

### Host prediction and host phylogenetic tree construction

We predicted the probable hosts of the rumen viruses using an alignment-dependent method (aligning prophage sequences and CRISPR spacer sequences with host genomes), which has a high prediction accuracy^80^. For prophage sequence alignment, we manually curated three databases: the databases of ruminant gut prokaryotes (including 22,095 bacterial MAGs and 410 archaeal MAGs from the same metagenomes collected for viral analysis in the current study), 2,729 rumen prokaryotic MAGs of the Cow Rumen genome database V1.0 (https://www.ebi.ac.uk/metagenomics/genome-catalogues/cow-rumen-v1-0), and 251,167 reference genomes of prokaryotes of NCBI RefSeq (release 211; downloaded in March 2022). The above prokaryotic MAGs and genomes were classified according to the GTDB taxonomy using GTDB-tk (option: -classify_wf)^81^. We aligned the representative vMAG sequence of each vOTU with the above prokaryotic genome/MAG sequences using BLASTn (option: -task metablast) to identify integrated prophage regions. A host match was called when >2,500 bp of a host genome or MAG matched a vOTU sequence at >90% sequence identity over 75% of the vOTU sequence length^6^. We predicted the probable protozoal hosts of the rumen viruses by searching the 52 high-quality SAGs^41^ for prophase regions using BLASTn and the above criteria.

We also predicted the probable hosts of the identified viruses by aligning the CRISPR spacer sequences of the vMAGs against the reference genomes and MAGs of prokaryotes. Briefly, we identified the CRISPR spacer sequences of the reference genomes and MAGs using MinCED (option: -minNR 2)^82^. The identified CRISPR spacer sequences were then aligned with the sequences of RVD using BLASTn (option: -dust no). A probable host was called when the CRISPR spacer sequence of a reference genome or MAG of prokaryotes matched exactly with a vMAG sequence (100% coverage and 100% identity). In total, we identified 43,166 vOTUs that had a CRISPR spacer match with their probable hosts or integrated into the host genomes. The sequences of these vOTUs were used to build a gene-sharing network using vConTACT2. After removing duplicated edges and clusters with <3 nodes, the network was imported into Cytoscape 3.7.2^83^ and annotated based on the taxonomy of the viruses and their hosts.

To display the infection patterns of rumen viruses, we constructed genus-level phylogenetic trees for the identified hosts of archaea, bacteria, and rumen ciliates. For the phylogenetic trees of bacterial and archaeal hosts, one genome was randomly chosen within each identified host genus. Then the 120 marker genes of bacteria and 122 marker genes of archaea of the genomes of the selected bacteria and archaea were aligned using GTDB-tk^81^. Then phylogenetic trees were constructed using the aligned marker genes and IQ-TREE (option: -redo -bb 1000 -m MFP -mset WAG,LG,JTT,Dayhoff -mrate E,I,G,I+G -mfreq FU -wbtl)^84^ and visualized using iTOL^79^. Lysogenic rates were calculated using VIBRANT. A ciliate tree was acquired from Li, et al. ^41^ and visualized also using iTOL^79^.

### Identification of auxiliary metabolism genes carried by vMAGs

We used stringent criteria to extract viral sequences, but during the initial manual curation of the rumen viral contigs, we found some contigs that were likely host genomic islands. Such contigs can be misidentified as viral sequences by virus identification tools^5^. Additionally, because it is still challenging to delineate the exact boundaries between host genomes and prophage genomes^35^ and any remaining host genes, which, if not removed, can be misidentified as AMGs, we performed a series of curations to select the vMAGs for AMG identification to minimize misidentification of host genes as AMGs. First, we only searched the complete vMAGs >10 kb (5,912 in total) for AMG identification using the criteria suggested in the benchmarking paper^34^. Then the vMAGs were further prepared for DRAMv^85^ annotation using VirSorter2 with the options ‘—prep-for-dramv’ applied. The complete vMAGs were then subjected to AMG identification and genome annotation using DRAMv. Second, the verified AMG-containing vMAGs in which an AMG was not flanked by both one of the viral hallmark genes and one viral-like gene or by two viral hallmark genes (category 1 and category 2 as determined by DRAMv) were removed. The AMG-containing vMAGs with AMGs at the end of vMAGs were also removed. Third, the remaining vMAGs were further manually curated based on the criteria specified in the VirSorter2 SOP (dx.doi.org/10.17504/protocols.io.bwm5pc86). We eventually obtained 1,880 vMAGs after the four steps of filtering. We manually checked the genomic context for these vMAGs and found that some of them were likely genomic islands. Therefore, we filtered the 1,880 vMAGs based on criteria established by Sun et al. (manuscript in preparation). Briefly, the vMAGs with only integrases/transposases, tail fiber genes, or any non-viral genes were removed. The remaining vMAGs were filtered again to remove those that did not have at least one of the viral structural genes (i.e., capsid protein, portal protein, phage coat protein, baseplate, head protein, tail protein, virion structural protein, and terminase) and the vMAGs containing genes encoding an endonuclease, plasmid stability protein, lipopolysaccharide biosynthesis enzyme, glycosyltransferase/glycosyl transferase families 11 and 25, nucleotidytransferase/nucleotidy transferase, carbohydrate kinase, nucleotide sugar epimerase. To benchmark our curation pipeline, 100 vMAGs were randomly selected from the 504 remaining vOTUs for detailed manual curation based on their genomic context. According to the benchmarking results, we were confident that we only retained vMAGs for AMG prediction. Detailed results of each curation step and full annotation of the final vMAGs and the annotation of the identified AMGs are in Table S4. Previously identified AMGs were identified from an expert-curated AMG database (https://github.com/WrightonLabCSU/DRAM/blob/master/data/amg_database.tsv).

### Identification of antimicrobial resistance genes carried by the rumen vMAGs

The vMAGs or contigs were searched for ARGs using the conservative criteria recommended previously^58^. Briefly, we downloaded the CARD database (v3.0.7)^56^ and searched it for ARGs carried by the vMAGs/contigs of RVD using BLASTp with a threshold of 40% alignment coverage and 80% sequence identity. We also searched for ARGs in RVD using the NCBI AMRfinder tools v.3.8.4^57^, which is a highly accurate tool and uses an expert-curated built-in database. The identified probable ARG-carrying vMAGs were curated to retain boda fida vMAGs using the pipeline to identify AMGs-containing vMAGs. To be conservative, the vMAGs with ARG at the end of the contig were removed unless it was adjacent to virus-related genes. A total of 22 ARG-carrying vMAGs were found, and they were manually checked individually to be viral origin based on genome context annotation. The detailed annotation and manual curation of individual ARG-carrying vMAGs are shown in Table S5. The representative vMAGs carrying each type of ARGs were picked based on CheckV completeness, and the number of cellular genes. The vMAGs with the highest completeness and least cellular genes were chosen as representative vMAGs and their genomic organizations were depicted individually using easyfig^86^ and annotated manually.

### Metagenomic read mapping and distribution of different viral populations

We estimated the abundance of ARG-carrying rumen viruses. Briefly, CoverM (option: -- min-read-percent-identity 0.95, --min-read-aligned-percent 0.75, --min-covered-fraction 0.7; https://github.com/wwood/CoverM) was used with the trimmed mean method, which calculates the average coverage after removing the regions with the top 5% highest and lowest coverage, to calculate the coverage of individual ARG-carrying vMAGs and the mapping rates of the metagenomic sequencing reads of 240 rumen metagenomes reported previously^36^ using RVD, RefSeq Viral, and IMG/VR. As high variability was seen in the read mapping rate results, we explored the results based on the metadata and compared the read mapping rate between healthy ruminants and those under SARA, and between RVD and IMG/VR with T-test using SciPy^87^. The obtained coverage was considered raw abundance, which was further normalized by library size to “coverage per gigabase of metagenome” as done previously^10^. We compared the normalized abundance of ARG-carrying vMAGs among different countries and different animal husbandry regimes using non-parametric Kruskal-Wallis test using SciPy^87^ with 100 bootstraps.

The core rumen virome was revealed based on the prevalence of vOTUs. First, the raw abundance table was transformed into a binary matrix (presence or absence) with a custom Python script. Then the prevalence of each vOTU in each sample was calculated. The vOTUs were categorized based on their prevalence in each species or type of ruminants as “individualized” (observed in only one sample), “rare” (observed in >1 but <20% of a specie or type), “common” (observed in 20 to 50% of a specie or type), and “core” (observed in >50% of a specie or type) vOTUs. The relative abundance of the “core” virome was calculated as the proportion of the “core” vMAGs to the combined abundance of vMAGs of a species or type. The presence of each “core” vOTU in each specie or type was plotted using the package ComplexHeatmap in R^88^.

---

**Table S1:**
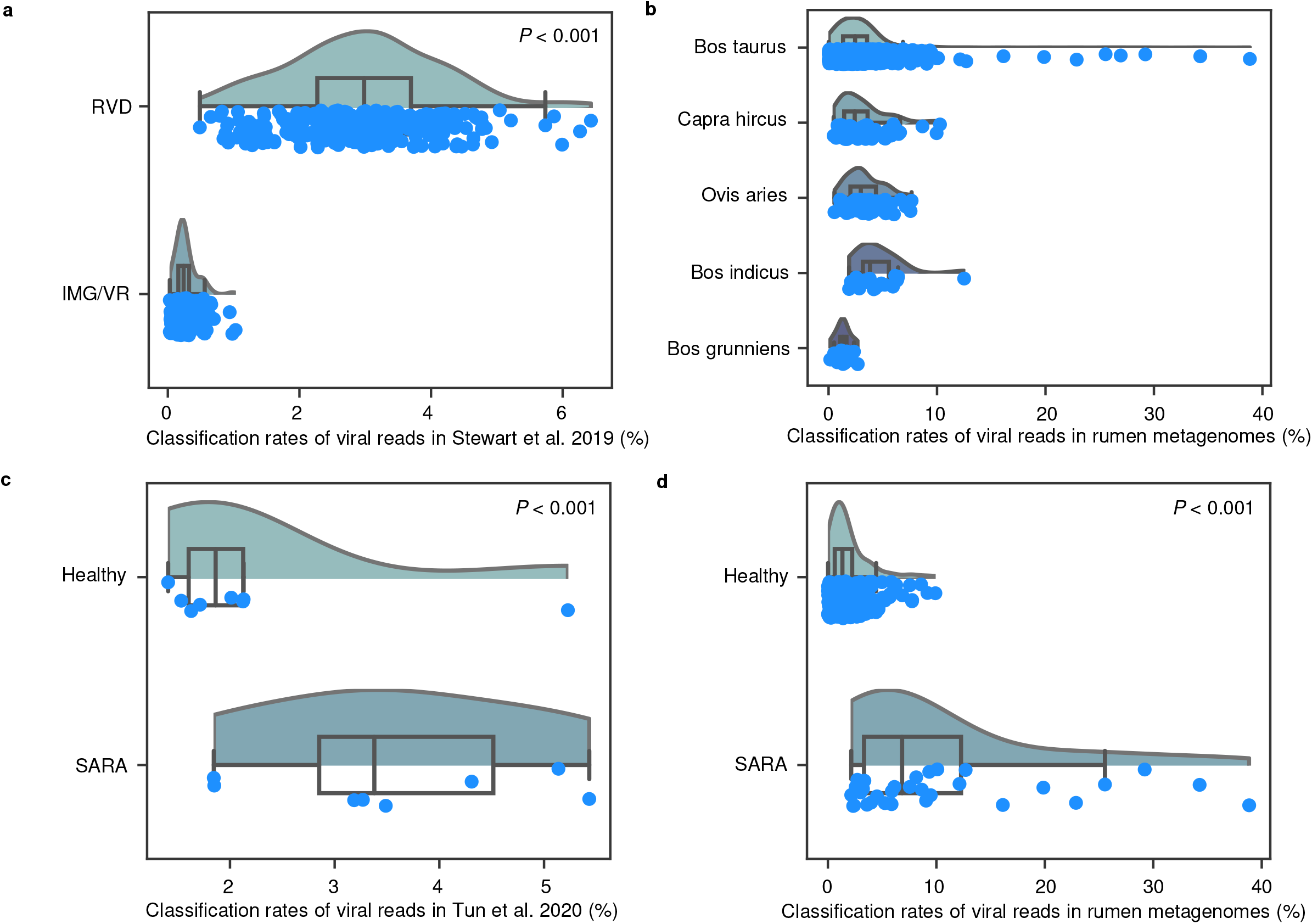
Mapping rates (%) of metagenomics sequence reads as viral sequences in metagenomes. **a**, Mapping rates of metagenomic sequencing reads across 240 rumen metagenomes reported by Stewart, et al. ^3^ with the IMG/VR and RVD. **b,** Mapping rates of metagenomic sequencing reads from Bos taurus (n=729), Capra hircus (n=82), Ovis aries (n=82), Bos indicus (n=23) and Bos grunniens (n=16). Ruminant species with less than 10 metagenomes were not included. **c,** Mapping rates of metagenomic sequencing reads from healthy goats (n=8) and goats under subacute rumen acidosis (SARA; n=8) reported by Tun, et al. ^4^. **d,** Mapping rates of metagenomic sequencing reads in healthy dairy cows (n=48) and dairy under SARA (n=203).

**Figure S2:**
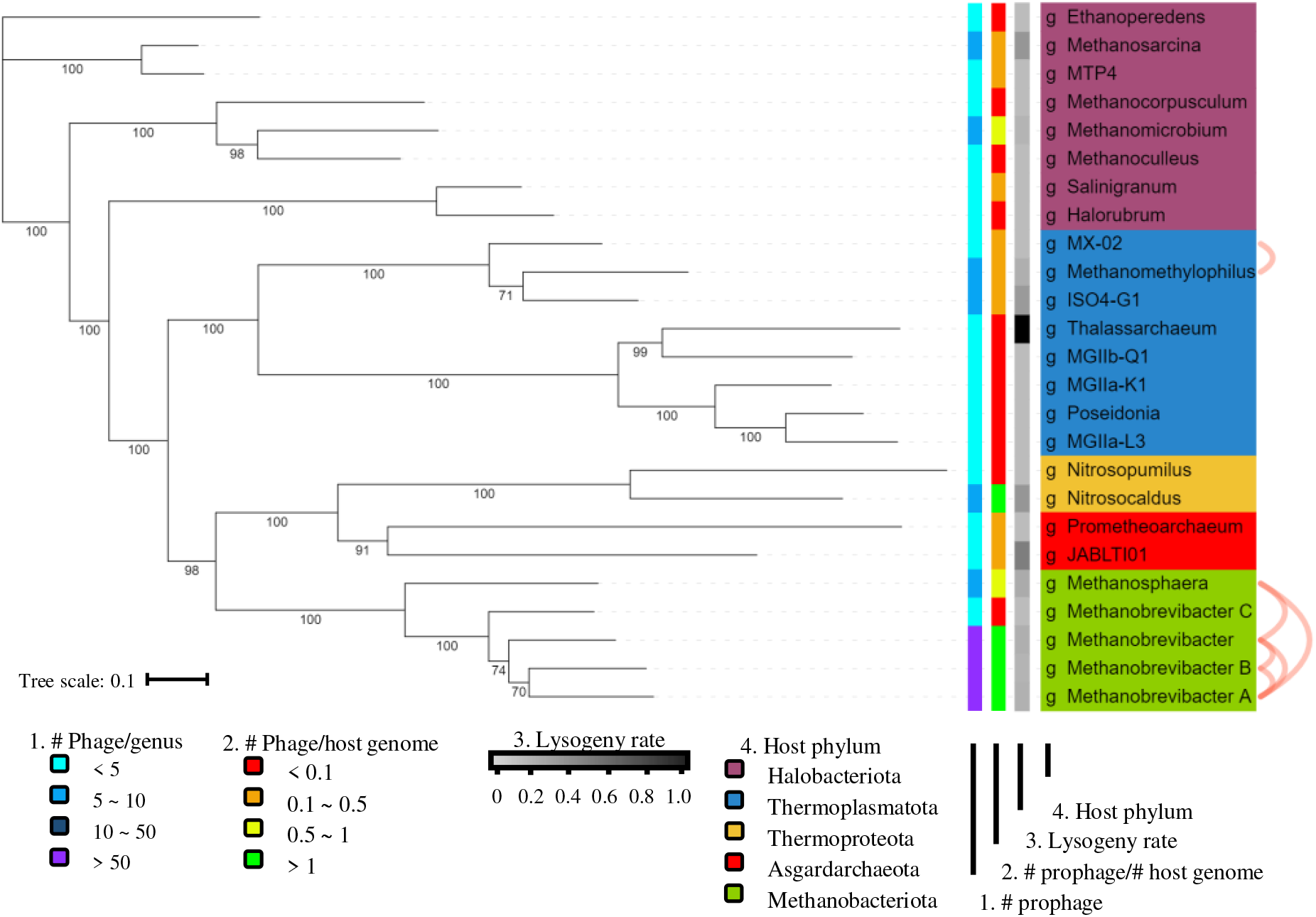
Predicted archaeal host of the rumen viruses. For vOTU with host across multiple genera, the predicted host genera were connected with orange arcs. The dendrogram represents the genome-based phylogenetic tree of 25 archaeal genera that contained the predicted hosts of 2,403 vOTUs. The hosts were inferred by (i) aligning the sequences of the representative vMAGs (the longest with the highest completeness) of each vOTUs with 410 metagenome-assembled genomes (MAGs) of rumen archaea and 8,367 bacteria reference genomes of NCBI RefSeq and (ii) aligning the CRISPR spacer sequences of the representative vMAGs and those of the RefSeq archaea genomes and MAGs. The RefSeq archaeal genomes and MAGs were classified using GTDB-Tk. The phylogenetic tree was constructed with the genomes or MAGs of the inferred hosts (clustered into genera) and their predicted phages to examine the lysogeny rate (calculated based on VIBRANT results), number of phages per genus, and number of phages per genome/MAG. In total, 6 vOTUs likely had a host range across multiple genera. These genera are connected with orange arcs.

**Figure S3:**
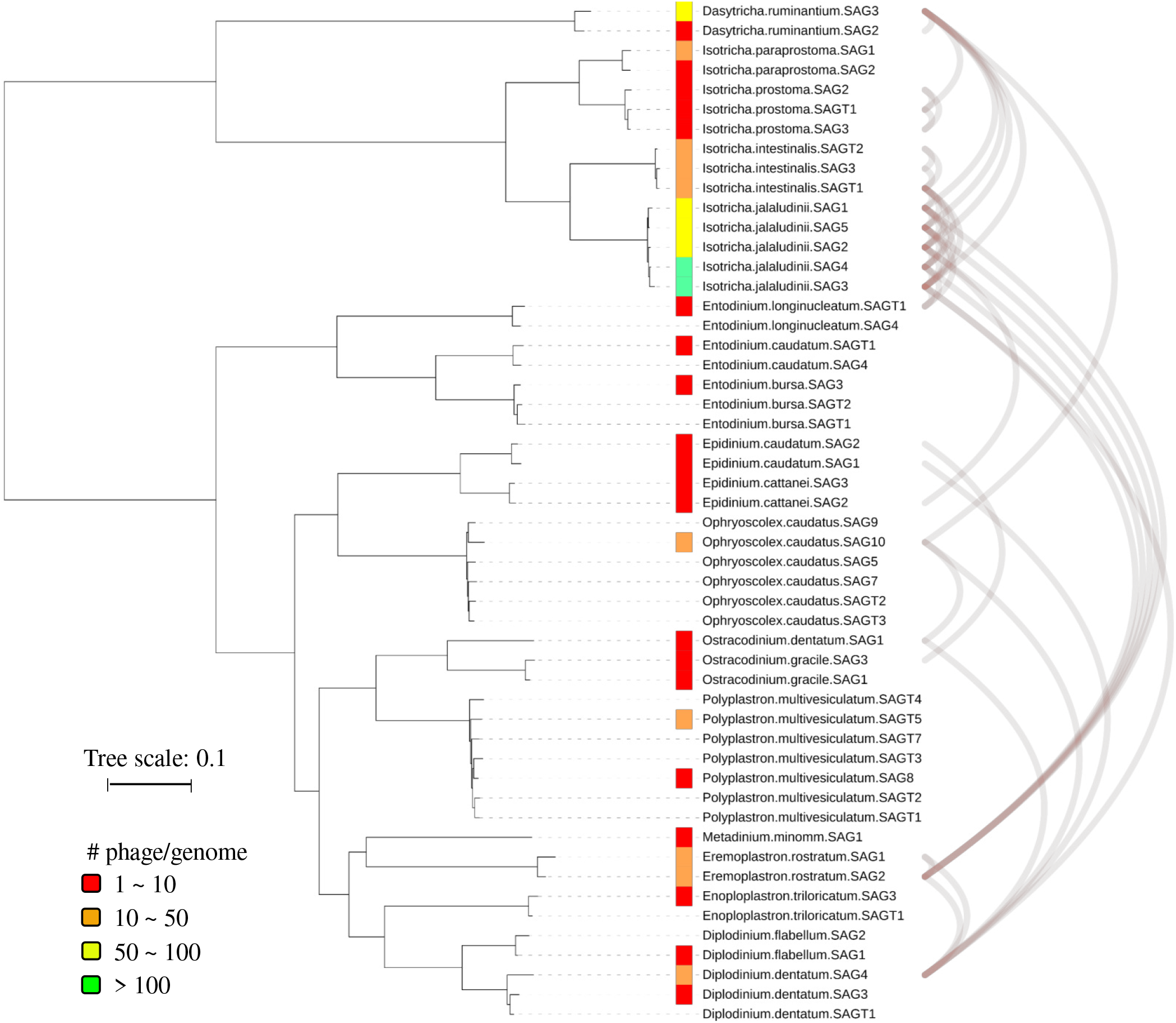
Predicted protozoal host of the rumen viruses. The host genomes were from the 52 high-quality ciliate genomes acquired from single amplified genome (SAG) and the dendrogram represents the phylogenetic tree of the 52 SAGs generated in the same study^5^. In total, 500 vOTUs were predicted to affect rumen ciliates. For vOTUs with host across multiple genomes, the host genome were connected with orange arcs. The phylogenetic tree was annotated according to the number of predicted prophages per SAG.

**Table S4:**
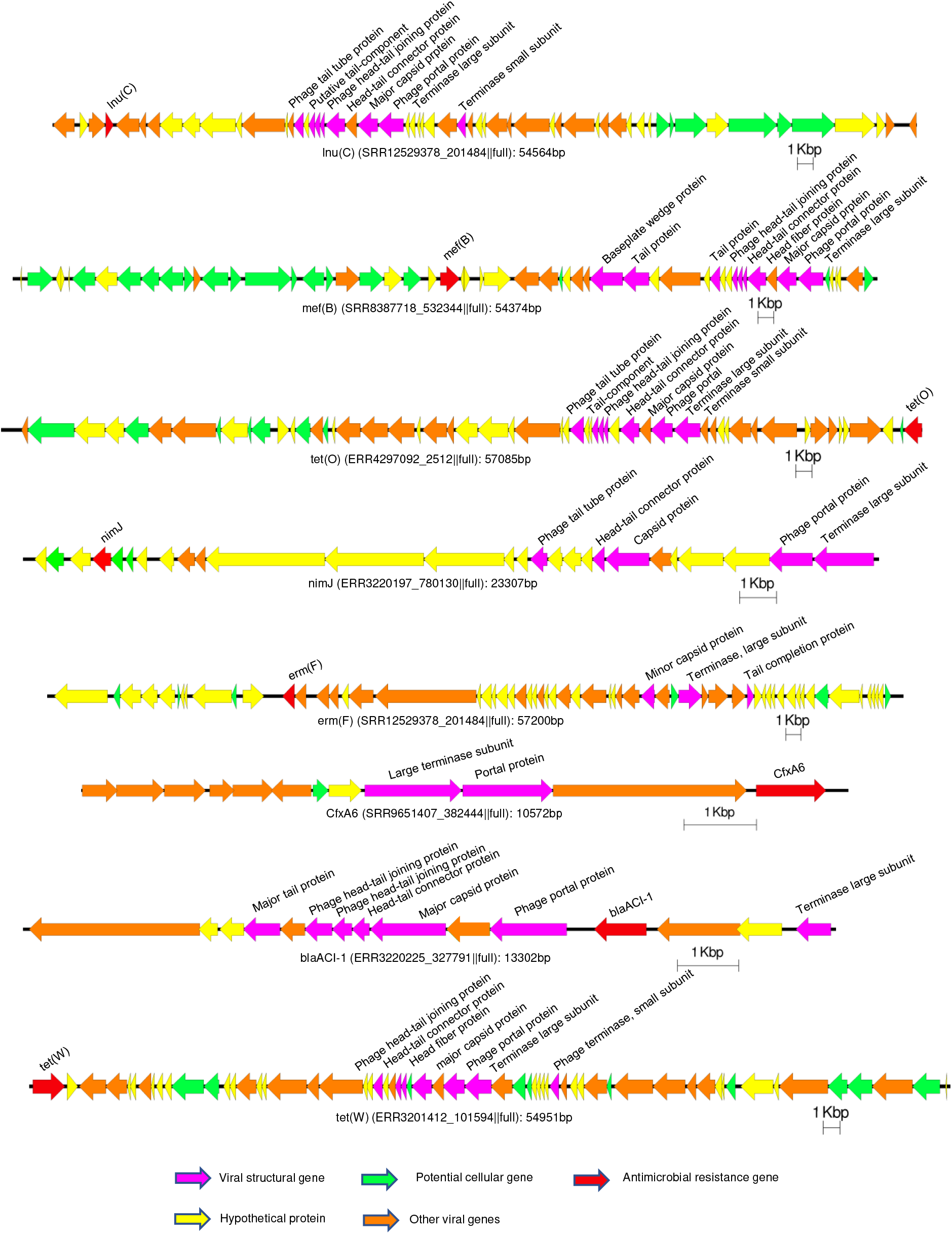
Genomic organization of representative viral contigs carrying each type of. **ARGs.** Representative viral contigs were chosen from each vOTU that had the highest completeness and least host genes.

## Notes

https://zenodo.org/record/7258071#.Y4aSnHbMK5c

### Competing Interest Statement

The authors have declared no competing interest.

